# Sample Size Impact (*SaSii*): an R script for estimating optimal sample sizes in population genetics and population genomics studies

**DOI:** 10.1101/2024.08.30.610119

**Authors:** Matheus Scaketti, Patricia Sanae Sujii, Alessandro Alves-Pereira, Kaiser Dias Schwarcz, Ana Flávia Francisconi, Matheus Sartori Moro, Kauanne Karolline Moreno Martins, Thiago Araujo de Jesus, Guilherme Brener Ferreira de Souza, Maria Imaculada Zucchi

**Author notes:** MS and PSS are joint first authors. PSS, KDS, AA-P and MIZ conceived the study. KDS and MS wrote and optimized the scripts. MS, TAJ and GBF performed the analyses. PSS, MS, AFF, MSM, KKMM and AA-P drafted the manuscript.

## Abstract

Obtaining large sample sizes for genetic studies can be challenging, time-consuming, and expensive. However, small sample sizes may generate biased or imprecise results. Many studies have suggested the minimum sample size necessary to obtain robust and reliable results, but it is not possible to define one ideal minimum sample size that fits all studies. Here, we aim to present SaSii (Sample Size Impact), a R script to help researchers to define minimum sample size and to indicate minimum sample size patterns for some taxa groups. The patterns were obtained by analyzing previously published datasets with SaSii and can be used as a starting point for the sample design of population genetics and genomic studies. Our results showed that it is possible to estimate an adequate sample size that accurately represents the real population and does not require the scientist to take time to write any program code, extract and sequence samples or use population genetics programs, making it easier to gather this information. We also confirmed that sample sizes of five to twenty-five for SNP and fifteen to thirty for SSR can be used for most plant species, giving a better direction for new studies.

## Introduction

Population genetics is an important and versatile tool to answer questions and to test hypotheses related to a broad range of research areas. The study of genetic patterns at the population level interconnects ecology and evolution, thus providing a framework to understand the impact of selection, genetic drift, gene flow on demography, and phenotypic frequencies [1]. Spatial-temporal approaches applied in landscape genetics can also be used to elaborate inferences of past population dynamics, human impacts in natural populations, and the effect of future environmental changes in populations [2]. In addition to population genetics contribution, genomic information is also becoming available and more accessible. Advances in genomic technologies reduced the price per data point, which is important to expand its impact to the development of both basic and applied research with model and non-model organisms [3].

All these applications and innovations require the development of methods to acquire and analyze data. Many publications over the last years focused on describing molecular marker choice [3], protocol refinement [4,5], and overall process decisions [6–8]. Another aspect that has been debated and is frequently questioned by journal reviewers is the minimum sample size. Poorly planned sample design and small sample sizes may prevent researchers to achieve robust results and conclusions. However, large sample sizes are often unfeasible due to time and budget reasons or for limitations associated with the species or populations characteristics, especially for endangered species.

Another challenge in defining sample size are differences in sampling guidelines. For microsatellite markers (SSR - Simple Sequence Repeats), the minimum sample size recommendation ranged from 20 to over 40 individuals [9,10]. For SNP (Single Nucleotide Polymorphism) data, recommended minimum sample size ranged from eight to over 50 individuals [5,11]. The variation is a result of differences in the hypothesis to be tested, objectives of the studies, and the taxa evaluated, so it is not possible to define one ideal minimum sample size that fits all studies [12].

When empirical data is not available before the study begins, simulation studies and extrapolation from similar taxa may be helpful in decision-making. All sampling guidelines cited above were defined by data simulation and rarefaction curves. These methods are already well known by statisticians, but writing the scripts to perform the analyzes may be difficult and time-consuming, which can be very discouraging to some researchers.

In this article, we aim to present *SaSii* (Sample Size Impact), a simulation program to help researchers to define and indicate minimum sample size patterns for taxa groups. The patterns discussed here were obtained by analyzing previously published datasets with *SaSii* and can be used as a starting point for the sample design of population genetics and genomic studies.

## Materials and Methods

### About *SaSii*

#### General idea of the script

We introduce a R [13] script designed to evaluate if the sample size is large enough to estimate robust genetic parameters from SSR and SNP data. The program reads the input dataset that contains individuals genotyped at a certain number of markers and estimates genetic parameters from subsamples of different sizes defined by the user. These estimates can be used to evaluate if the sample size is large enough to adequately represent the genetic diversity observed in the original complete dataset. The script, tutorials and other documentation are available at https://sasii.readthedocs.io/en/latest/index.html.

#### Data input and config file

The program can read both SSR and SNP markers of multi locus and diploid individuals’ data, formatted as Structure [14] data file, accepting some minor variations of the original format, and saved as comma-separated values file (.csv). The individuals’ data are coded in rows and loci in columns. Alleles must be represented as numbers, with missing data coded as 0 (zero) or −9. The user may use two types of data organization, one in which each individual is in one row and genotypes in two consecutive columns (the two alleles of the same locus are in two consecutive columns - Fig S1), and another in which each individual is in two consecutive rows and each locus in one column (the two alleles of the same locus are in two consecutive rows - Fig S2).

In order to manage data information of the command-line version, the user needs to fill a configuration file (hereafter *config* file) with parameters used to describe the data and the file structure, such as file name, input file organization, missing data code, line number with the first individual of the population. The user also must define in this *config* file the following analysis settings: size of the smallest subsample, number of resamples of each subsample size and minimal frequency of allele to be preserved.

### Random subsampling of the dataset

The program makes random subsamples of the input dataset without replacement according to the settings defined by the user in the *config* file. For example, with a simulated dataset consisting of 50 individuals, with the smallest subsample size of 5, the program will create new datasets with the following subsample sizes: 5, 10, 15, 20, 25, 30, 35, 40, 45, and 50. The smallest subsample size also defines the intervals between them.

For each subsample, the program estimates genetic parameters and compares them to the original input dataset (Fig S3). It also calculates average and standard deviations from the resamples of each subsample size and compares its estimates with the original dataset estimates to create the output tables and plots. The parameters estimated are: allele frequencies, observed heterozygosity (*H_O_*), expected heterozygosity under Hardy-Weinberg Equilibrium (*H_E_*), and genetic distances between the original dataset and the subsample (Wrigth’s *F_ST_*, Nei’s Distance and Rogers’ distance).

### Data output and sample size impact analysis

To estimate the minimum sample size, we iterate over all individuals at each sample size, detecting alleles and calculating their frequency. The program outputs 12 pairs of files, each pair consists of one numerical file and one plot with the results of the genetic parameter estimates, which are described briefly below.

### Fraction of common alleles detected (output 1)

For some analysis, common alleles (frequency > 0.95) are more informative [10]. *SaSii* identifies the most common alleles in the input dataset and calculates their percentages in each subsample.

The output 1 has a boxplot of the fraction of common alleles that are represented in the subsamples of each size (Fig 1A, E). The red line in the plot indicates the threshold above which 95% of the common alleles are present in the subsamples.

**Figure 1.**
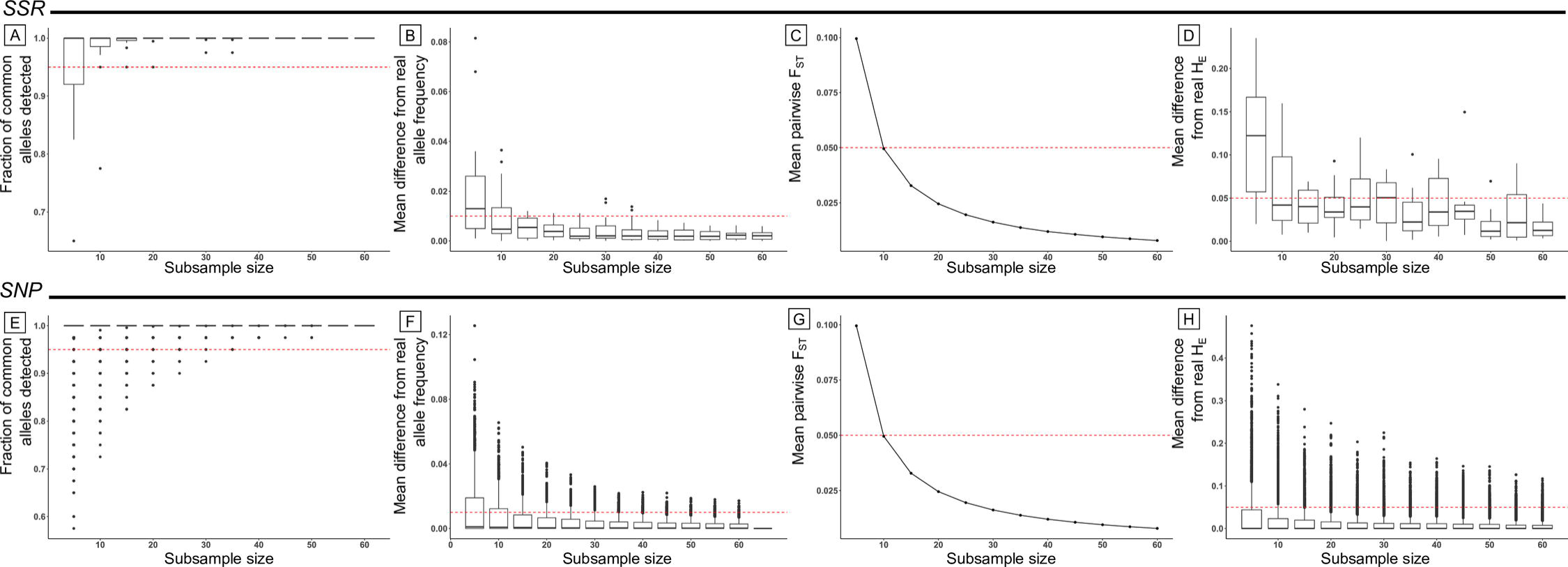
Results of *SaSii* analyzes for SSR (top) and SNP (bottom) simulated data. Only the smallest subsample sizes are show. A, E: Amount of individuals with more than 95% of alleles detected; B,F: mean difference of allele frequencies between the sampled populations and the original dataset; C, G: mean pairwise *F_ST_* between the original and the simulated datasets; D, H: mean of expected heterozygosity of populations within a same class (same population and sample size).

### Mean difference from original allele frequency (output 2)

The program calculates the difference between the allele frequencies of the original dataset and the subsamples. The table output has the average difference for each allele for every subsample size. The output plot has a boxplot of these differences for each subsample size (Fig 1B, F). The red line indicates the threshold below which there is a 0.01 mean difference in allele frequencies from the original dataset and the subsample.

### Frequencies of the most and least common alleles (output 3)

*SaSii* identifies the most and the least common alleles overall loci in the original dataset and estimates their frequencies in each subsample. The output table contains the mean allele frequencies, the standard deviations, the minimum and the maximum allele frequencies over subsamples. In addition, these results are also plotted in a boxplot graph, which contains a vertical line with the minimum and maximum values, a bold central line with the mean value and a box with the standard deviations.

### Expected heterozygosity (outputs 4 and 5)

Genetic diversity often requires an accurate estimation of *H_E_* in the real population. So, for each sample size, *SaSii* calculates *H_E_* (per locus in output 4 and averaged overall loci in output 5) of the simulated subsample to compare their genetic diversity in relation to the original dataset. In the tables of outputs 4 and 5, the user can find the mean *H_E_* for each subsample size, the standard deviation over subsamples, the minimum and maximum estimates over the subsamples.

The output 4 plot is a boxplot of estimates for each subsample size. The output 5 plot contains a vertical line with the minimum and maximum *H_E_* values, a bold central line with the mean value and a box with the standard deviations.

### Observed heterozygosity (outputs 9 and 10)

Outputs 9 and 10 are equivalent to outputs 4 and 5 but contain *H_O_* estimates. For each sample size, *SaSii* calculates *H_O_* (per locus in output 9 and averaged overall loci in output 10) of the simulated subsample to compare genetic diversity in relation to the original dataset. In the tables of outputs 9 and 10, the user can find the mean *H_O_* for each subsample size, the standard deviation over subsamples, the minimum and maximum estimates over the subsamples.

The output 9 plot is a boxplot of estimates for each subsample size. The output 10 plot contains a vertical line with the minimum and maximum *H_O_* values, a bold central line with the mean value and a box with the standard deviations.

### Mean pairwise F_ST_ and genetic distances (outputs 6, 7 and 8)

*SaSii* calculates the genetic differentiation (*F_ST_* [15]), Nei’s distance [16], and Rogers’ distance [17] between subsamples and the original dataset to check if the subsamples are giving accurate and consistent results which may be indicated by small genetic distances [10]. Mean values over loci and standard deviation are estimated and presented in the output tables. The output plots show the average values over subsamples for each subsample size. The *F_ST_* plot has a red line with the 0.05 threshold, that indicates low population genetic differentiation (Fig 1C, G).

### Mean difference from the original heterozygosities (outputs 11 and 12)

Outputs 11 and 12 show the differences between *H_O_* and *H_E_* (respectively) from the original dataset and the subsamples. The output tables contain the mean difference over subsamples for each locus and subsample size. The output plots contain boxplots of the mean differences in heterozygosities over loci and subsamples for each subsample size. Figs 1D and H show the plots for *H_E_*. The red line in the plots indicates greater deviances than 0.05.

### Sample size impact analysis

To estimate the minimum sample size, we suggest that the users analyze the predefined threshold for each plot. All sample sizes that pass these thresholds can be considered large enough for that parameter. When different parameters indicated different minimum sample sizes, we suggest a conservative approach, selecting the largest size as the overall minimum. For a better analysis we suggest four graphs: outputs 1, 2, 6, and 11, that summarizes most of the genetics parameters for the species.

### Script validation

In order to evaluate *SaSii*’s predictive accuracy, we constructed two datasets using EasyPop 2.0.1, a forward simulator, to generate similar data to those found in nature [18]. These populations were intended to simulate SSR and SNP markers, using the following parameters: ploidy (2), sex number (2), mating system (Random), number of populations (1), population size (600), male and female ratio (1:1), free recombination (yes), same mutation scheme (yes), mutation model (mixed model of SSM with a proportion of KAM mutation events), proportion of KAM events (0.5), variability of initial population (maximal) and number of generations (10). The number of loci, mutation rate and number of possible allelic states were different for SSR and SNP markers, respectively: 10 and 2,500 loci, 1 × 10^-4^ and 1 × 10^-7^ for mutation rates and 20 and two allelic states.

Then, we configured SaSii’s input setting the default value for all common parameters (resampling number, sample size, minimum frequency and max sample size) for both simulated datasets. SaSii generated fifty subsamples from the amount of each population’s size, with a sample size equals to five, selecting allele frequency with at least 0.05. The remaining parameters were defined depending on the data file of each species. All datasets were analyzed in a workstation with the following configuration: AMD Ryzen 7 5700X, 32GB RAM.

### Patterns to predict the minimum sample size

We investigated if any characteristics of the species or the type of molecular marker could be used as a predictor for generalizations about the minimum sample size necessary for genetic estimates. We used empirical datasets from population genetic studies: one insect, one reptile, and 15 plant populations, of which seven were SSR and eight were SNP data as shown in Table 1. The results found here for each species may be applicable for other studies and improve the decision-making process in population genetic studies. *SaSii* runs for all species followed the same settings as the validation analyses.

**Table 1.** Characteristics of the datasets.

| Group | Marker<br>type | Species | Mating<br>system | Sample<br>size | Number<br>of loci | Reference |
| --- | --- | --- | --- | --- | --- | --- |
| Plant | SNP | *Acrocomia aculeata* | Mixed | 25 | 3,269 | [19] |
|  |  | *Casearia sylvestris* | Allogamy | 23 | 1,257 | [20] |
|  |  | *Chloropyron maritimum* | Allogamy | 15 | 111 | [21] |
|  |  | *Euterpe precatoria* | Allogamy | 12 | 3,803 | Francisconi et al. (work is in progress) |
|  |  | *Manihot dulcis* (sweet manioc) | Allogamy | 46 | 1,985 | [22] |
|  |  | *Manihot esculenta* (bitter manioc) | Allogamy | 53 | 1,985 | [22] |
|  | SSR | *Bertholletia excelsa* | Allogamy | 91 | 12 | [23] |
|  |  | *Caryocar villosum* | Allogamy | 38 | 7 | [24] |
|  |  | *Copaifera langsdorffii* | Mixed | 357 | 8 | [25] |
|  |  | *Erythrophleum suaveolens* (Camaron) | Mixed | 177 | 9 | [26] |
|  |  | *Erythrophleum suaveolens* (Democratic Republic of Congo) | Mixed | 88 | 9 | [26] |
|  |  | *Eugenia dysenterica* | Allogamy | 28 | 7 | [27] |
|  |  | *Metrosideros polymorpha* | Allogamy | 319 | 9 | [28] |
|  |  | *Myroxylon peruiferum* | Mixed | 50 | 9 | [29] |
|  |  | *Shorea macrophylla* | Mixed | 88 | 14 | [30] |
|  |  | *Solanum lycocarpum* | Allogamy | 178 | 5 | [31] |
| Animal | SNP | *Tetragonisca angustula* | - | 32 | 3,573 | [32] |
|  | SSR | *Caiman crocodilus* | - | 25 | 11 | [33] |

Information used to evaluate patterns for minimum sample size predictors.

Here, the minimum sample size was defined based on the proportion of common alleles detected, mean difference of allele frequencies, mean pairwise *F_ST_*, and mean difference in *H_E_*. We tested the null hypothesis of no differences in minimum sample size for datasets of SNP and SSR markers using all plant and animal datasets. We also tested the null hypothesis of no difference in minimum sample size for mating systems in allogamous and mixed mating plant datasets. Furthermore, we used a two-sided Wilcoxon rank sum test in R [13] to perform the statistical test.

## Results

### Script validation

Considering the simulated datasets with six hundred samples, the mean *H_E_* was 0.9468 for SSR and 0.4967 for SNP. In Fig 1, we show the graphs generated by SaSii used to infer the minimum sample size. For the SSR data, the minimum sample size ranged from ten to forty-five, depending on the parameter analyzed (Figs 1A, D). In Fig 1A and C, it was noted that for subsamples with a minimum of ten individuals, at least 95% of common alleles were detected and the mean *F_ST_* was smaller than 0.05. However, in the graphs that presented the differences in the original allele frequencies and *H_E_* (Figs 1B and D), we noticed the minimum individual number was at least fifteen and forty-five individuals. With populations with higher individual numbers, the boxplot interval did not have a relevant increase of precision.

Following the same strategy for the SNP data, the graphs suggested that five individuals were necessary to have a minimum of 95% of alleles detected (Fig 2E), and a low mean difference in *H_E_* (Fig 2H), while ten and fifteen individuals were sufficient for a *F_ST_* smaller than 0.05 (Fig 2G), and a low mean difference allele frequency (Fig 2F).

**Figure 2.**
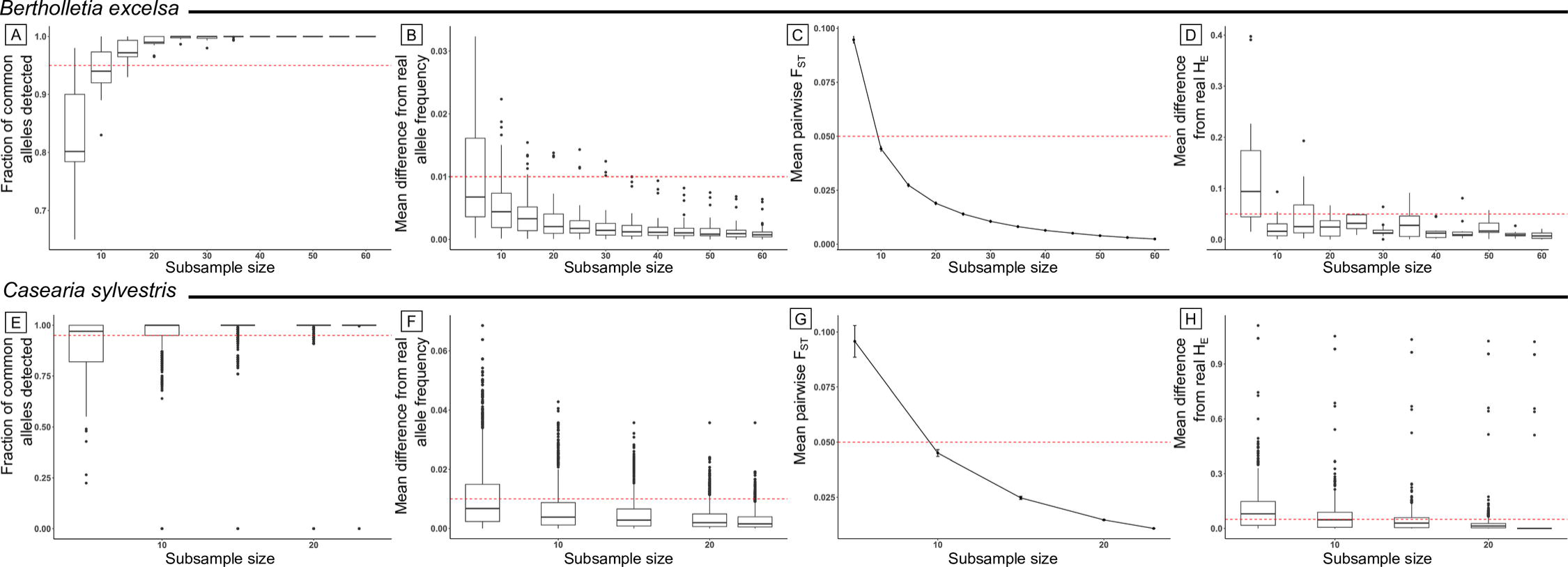
Results of *SaSii* analyzes for *Bertholletia excelsa* and *Casearia sylvestris* datasets. A,E: proportion of common alleles detected in each subsample; B,F: mean difference of allele frequency between the sampled populations and the empirical dataset; C,G: mean pairwise *F_ST_* between subsample and complete dataset; D,H: mean difference in *H_E_* estimates between subsamples and the complete dataset.

### Patterns to predict the minimum sample size

We analyzed data from species of plants and animals and observed minimum sample size ranging from five to 25 (Table 2). All results files can be found in https://sasii.readthedocs.io/en/latest/index.html. For each species, we observed that the minimum sample size could be different depending on the parameter analyzed.

**Table 2.**
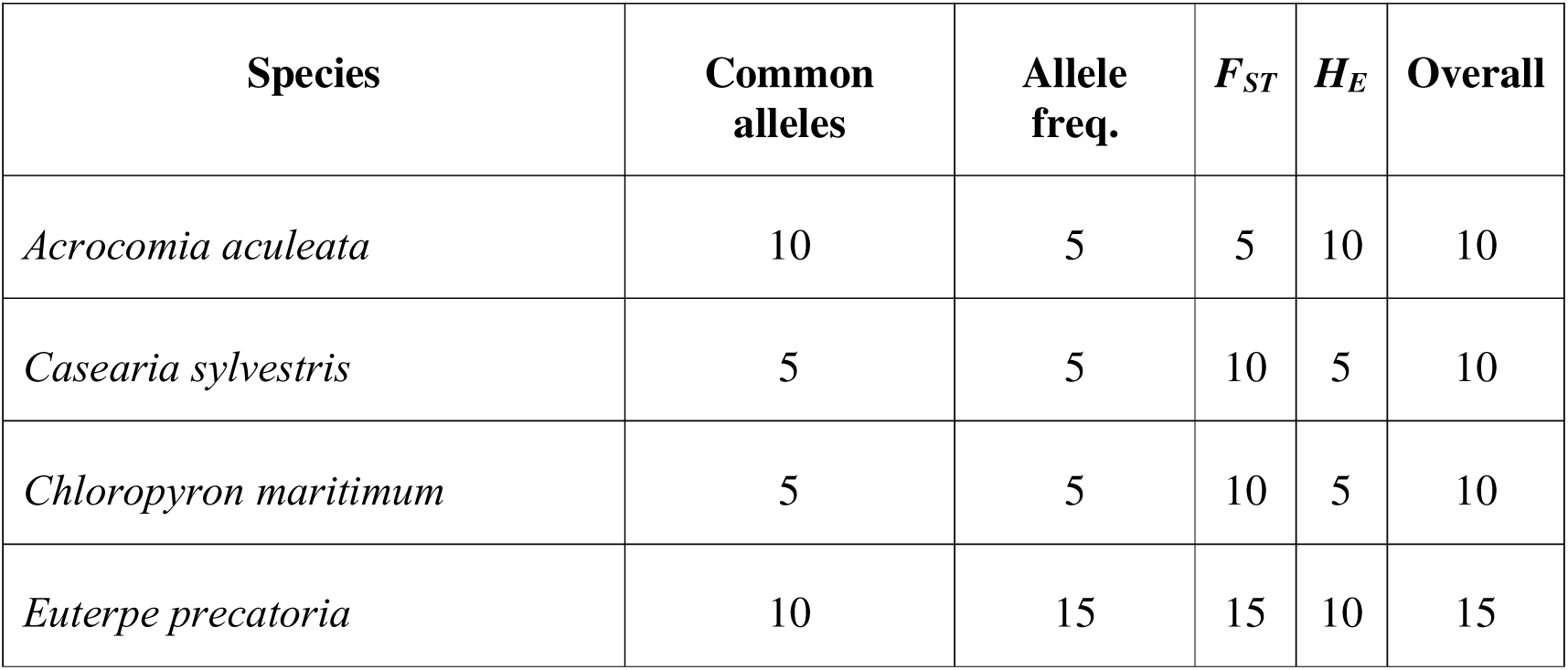
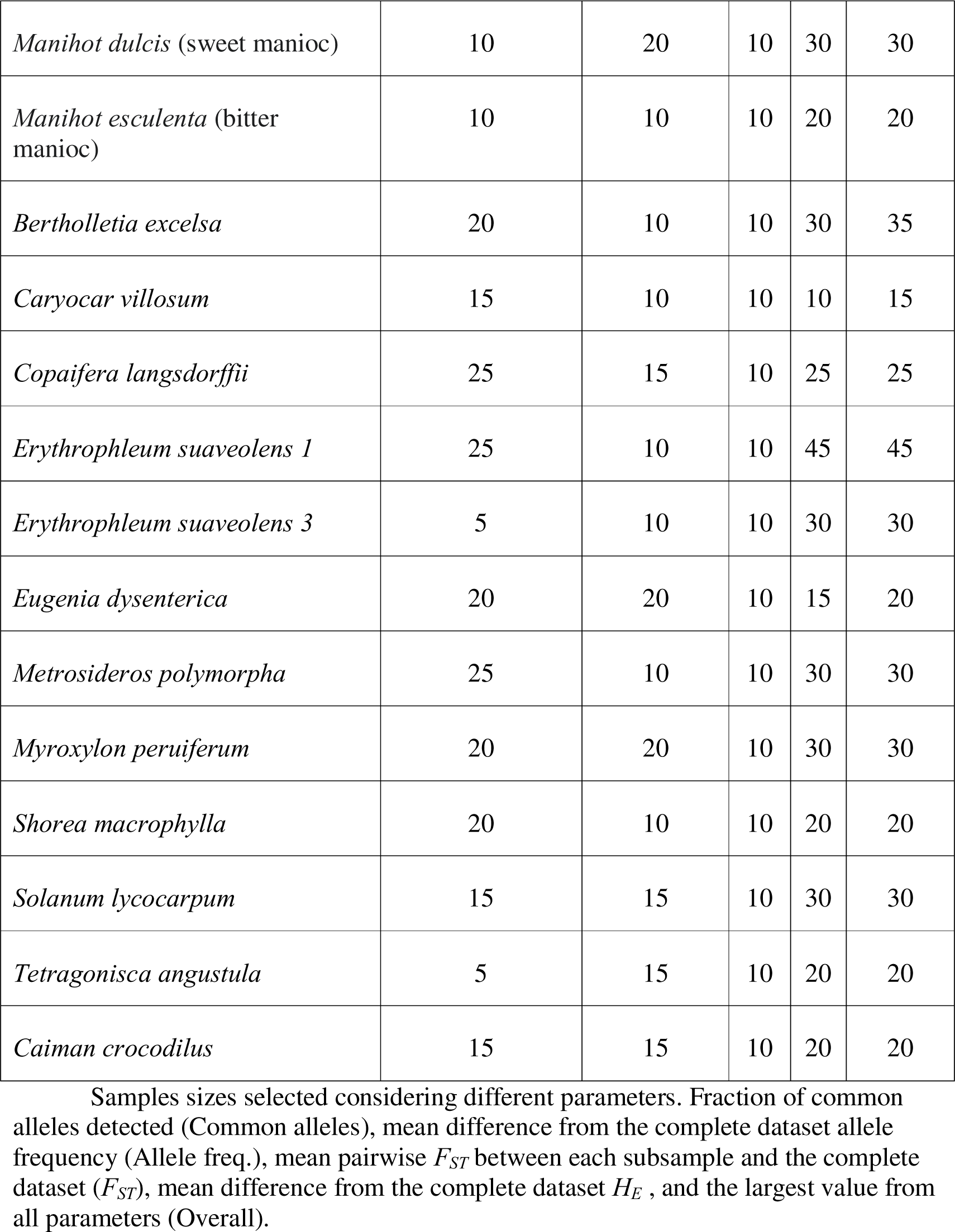
Minimum sample sizes detected with *SaSii*.

| Species | Common alleles | Allele freq. | $F_{ST}$ | $H_E$ | Overall |
| --- | --- | --- | --- | --- | --- |
| *Acrocomia aculeata* | 10 | 5 | 5 | 10 | 10 |
| *Casearia sylvestris* | 5 | 5 | 10 | 5 | 10 |
| *Chloropyron maritimum* | 5 | 5 | 10 | 5 | 10 |
| *Euterpe precatoria* | 10 | 15 | 15 | 10 | 15 |
| *Manihot dulcis* (sweet manioc) | 10 | 20 | 10 | 30 | 30 |
| *Manihot esculenta* (bitter manioc) | 10 | 10 | 10 | 20 | 20 |
| *Bertholletia excelsa* | 20 | 10 | 10 | 30 | 35 |
| *Caryocar villosum* | 15 | 10 | 10 | 10 | 15 |
| *Copaifera langsdorffii* | 25 | 15 | 10 | 25 | 25 |
| *Erythrophleum suaveolens 1* | 25 | 10 | 10 | 45 | 45 |
| *Erythrophleum suaveolens 3* | 5 | 10 | 10 | 30 | 30 |
| *Eugenia dysenterica* | 20 | 20 | 10 | 15 | 20 |
| *Metrosideros polymorpha* | 25 | 10 | 10 | 30 | 30 |
| *Myroxylon peruiferum* | 20 | 20 | 10 | 30 | 30 |
| *Shorea macrophylla* | 20 | 10 | 10 | 20 | 20 |
| *Solanum lycocarpum* | 15 | 15 | 10 | 30 | 30 |
| *Tetragonisca angustula* | 5 | 15 | 10 | 20 | 20 |
| *Caiman crocodilus* | 15 | 15 | 10 | 20 | 20 |

Samples sizes selected considering different parameters. Fraction of common alleles detected (Common alleles), mean difference from the complete dataset allele frequency (Allele freq.), mean pairwise $F_{ST}$ between each subsample and the complete dataset ( $F_{ST}$ ), mean difference from the complete dataset $H_E$ , and the largest value from all parameters (Overall).

For example, we identified that for *Bertholletia excelsa*, a species evaluated with SSR markers (91 samples, 12 loci), 15 individuals were sufficient to detect at least 95% of the common alleles and small (< 0.05; Fig 2A). Subsamples of 10 or more individuals presented *F_ST_* values smaller than 0.05, mean differences from the original *H_E_* and mean allele frequencies were sufficient to achieve a small difference when compared with the original dataset (N = 91; Fig 2B, C, D).

For SNP data, the minimum sample size was smaller than SSR. We show the analysis of *Casearia sylvestris* (23 samples, 1,257 loci) as an example. For this species, with only five individuals we obtained a good representation of the original data, as shown by the fraction of common alleles detected and the mean difference of allele frequencies (Figs 2E and F). However, subsamples with less than 10 individuals showed higher *F_ST_*and larger mean *H_E_* differences from the original dataset (Figs 2G and H), suggesting the need of additional individuals for more robust results. Considering this, the minimum sample size may range between 5 and 15 individuals, depending on the criterion adopted and the objectives of the study.

Comparing other SSR marker data, we could identify that for the detection of common alleles, most of the species showed that at least fifteen individuals were necessary for sufficient representation. *C. langsdorffii*, *M. peruiferum* and *C. crocodilus* needed more individuals. As for other graphs, most species had the same sample size considering the different parameters, with five individuals for differences in allele frequencies and ten for the mean difference of *F_ST_* and mean *H_E_*. The overall minimum sample size found for SSR markers was between fifteen and twenty-five individuals. For SNP datasets, the majority overall minimum sample size was ten individuals, being influenced by the *F_ST_* estimated between the complete dataset and each subsample. Only *T. angustula* and *C. sylvestris* needed twenty and fifteen individuals to better represent the original dataset (i.e. small mean differences from original *H_E_* and small *F_ST_*). For the detection of common alleles and allele frequencies, usually only five individuals were needed.

We observed that minimum sample sizes were smaller for SNP than for SSR dataset (*p* = 0.018), Fig 3 left. However, there was no significant difference (*p* = 0.471) between minimum sample sizes for plants with different mating systems (Fig 3 right).

**Figure 3.**
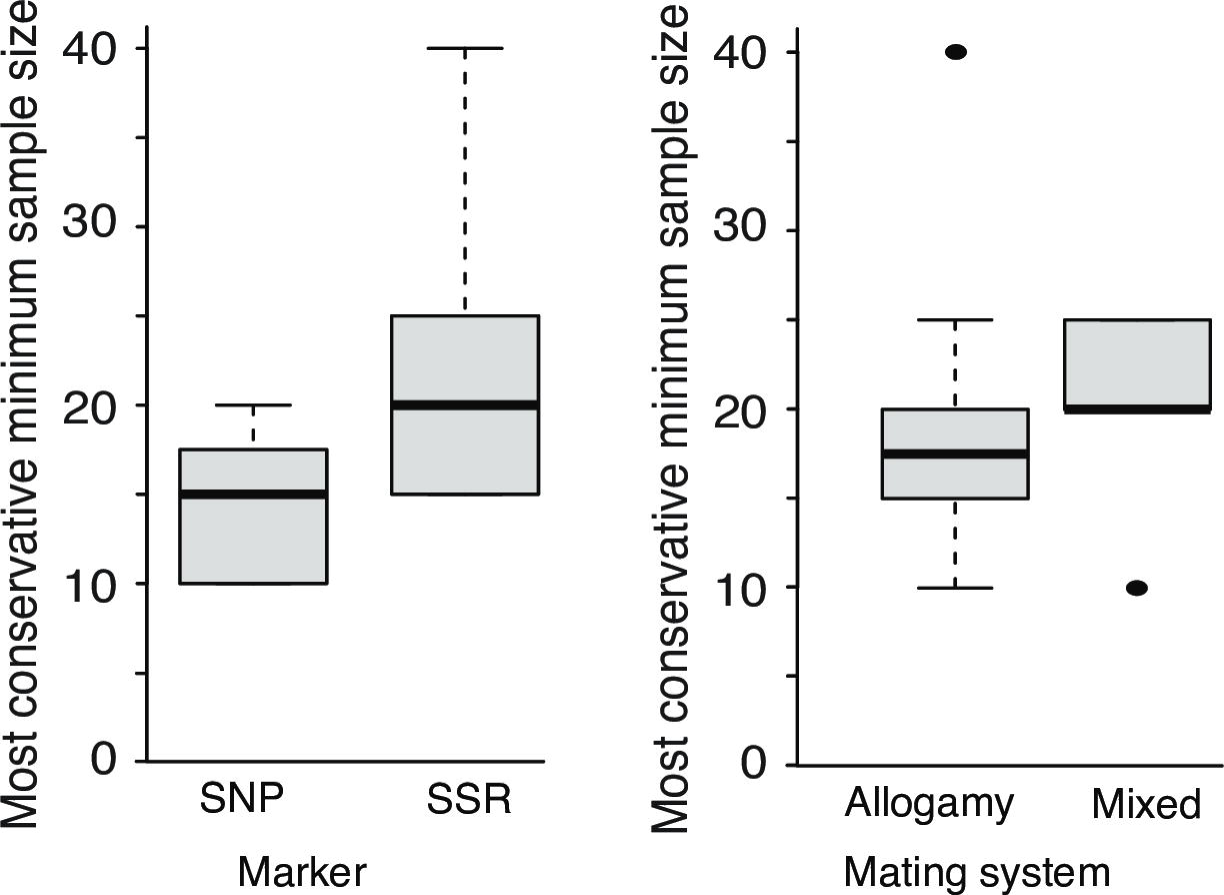
Minimum overall sample sizes observed for empirical datasets grouped by molecular marker type (left) and mating system (right).

## Discussion

*SaSii* script was efficient to analyze both SSR and SNP datasets, using computers with standard configurations. Our results showed that the minimum sample sizes varied between analyzed species and depended on the parameter evaluated. Overall, for SNP data, a smaller sample size was necessary to obtain robust results when compared to SSR data results. The script presented here can be used both to help developing sample strategies and to evaluate if the samples collected are sufficient for the analyses.

The analysis of simulated data indicated that SaSii can provide precise estimations of *H_O_* and *H_E_*. Larger subsamples show values for these parameters similar to the total dataset and smaller subsamples show higher variance in estimates. For SSR simulated data, twenty-five individuals were enough to represent the entire dataset. For SNP data, the minimum sample size was ten individuals. It is well known that the sample size for SSR and SNP data are around twenty-five to forty and eight to fifty, respectively [5,9,10,11]. This number may vary depending on the species, location, and study focus. Our simulation results confirm these recommendations.

As for our empirical SSR datasets, we found that, for some populations and species, the minimum sample size was smaller than twenty-five to forty individuals indicated in the literature [9,10]. For empirical SNP data, our results were similar to the literature suggestions [5,11]. Our results indicated that for species analyzed with SSR markers, the sample size should be around fifteen to twenty-five and, while for SNPs, the recommended sample was between five and twenty. This highlights the importance of taking the particularities of each species into account. When using *SaSii* to help to determine sampling sizes, we suggest that the researchers evaluate the *F_ST_* and *H_E_* parameters for SNP data, and the fraction of common alleles detected and *H_E_*for SSR data.

Our results suggest that most SSR and SNP data usually presented similar sample size requirements, within marker’s type. However, some species need larger sample sizes, as reported for the species *C. langsdorffii, M. peruiferum* and *C. crocodilus* (SSR) and *T. angustula* and *C. sylvestris* (SNP). Firstly, *C. crocodilus* and *T. angustula* differ from the other because they are both animal species, which can have distinct characteristics that can impact the minimum sample size. Beyond this, in plants, factors such as pollination syndrome, seed dispersal and population characteristics (for example, density, genetic structure, allelic richness, effective population size) can be influencing the minimum sampling size [11]. *For C. langsdorffii*, elevated levels of inbreeding, which resulted in small estimates of observed heterozygosity, were reported in different genetic studies of natural populations [34,35]. The *empirical* dataset for this species may be reflecting this condition, which resulted in a larger sample size than for the other species that used the same type of marker.

After getting *SaSii* outputs, we evaluated if the marker type and the mating system could influence the minimum sample size. This turned out to be true for marker type: as suggested by previous studies, when using SNP markers lower sampling sizes than SSR markers would be required to represent the species’ genetic diversity. However, we did not find a significant difference between minimum sample size in plants with different mating systems. We believe that *SaSii* can aid future genetic researchers providing tools for simulating populations and plotting graphs that summarize the most important parameters for population genetics studies. From our simulated and empirical data output, we propose that it is possible to reduce even more the sample sizes from SSR and SNP data, cheapening genetics studies costs.

Also, considering the number of SNP produced in recent articles, we recommend that *SaSii* should be used for analyzing genomic data with up to 15,000 SNPs and with at most three hundred individuals, our tests showed a limitation on memory usage for R in datasets with higher SNP or individuals’ numbers. It is possible to use larger datasets, for this, the user should have access to a better workstation, however, with the disadvantage of a higher time consumption.

## Conclusion

We presented *SaSii* a R script that takes SNP and SSR data to calculate estimates of genetic diversity and provide plots as outputs to help researchers deciding a reasonable sampling effort for their genetic studies. Our results showed that it is possible to estimate an adequate sample size that represents populations and do not require that the scientist takes time to write any program code, extract and sequence samples or use genetics population programs, making it easier to gather this information. Our results showed that a sample size per population of 5 to 25 for SNP and 15 to 30 for SSR could be used for most plant species, giving a better direction for new studies.

## Acknowledgements

We thank Chico Mendes Institute for Biodiversity Conservation (ICMBio) and SISGEN for sampling and research permits. This study was supported by various grants, including São Paulo Research Foundation (FAPESP 2018/00036-9, 2013/00003C0 and 2012/08307C5, 2011/50296-8, 2011/21295-3, 2012/03246-8, 2012/06431-0, 2013/05762-6, 2013/08086-1, 2013/11137-7, 2014/01364-9, 2015/06349-0), and and the National Council for Scientific and Technological Development (CNPq 313417/2023-7)

## Supporting Information

**S1 Fig.**
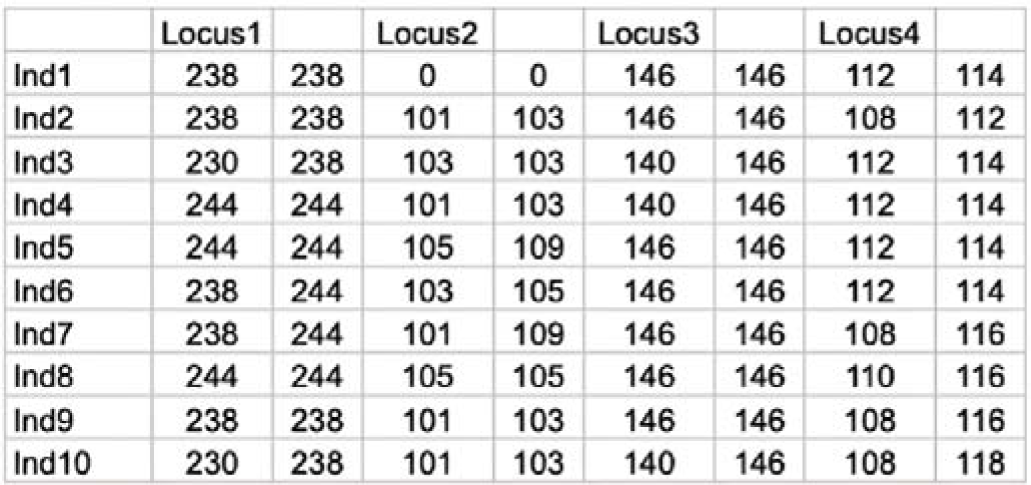
Data example for datasets with the two alleles of the same locus are in two consecutive columns.

**S2 Fig.**
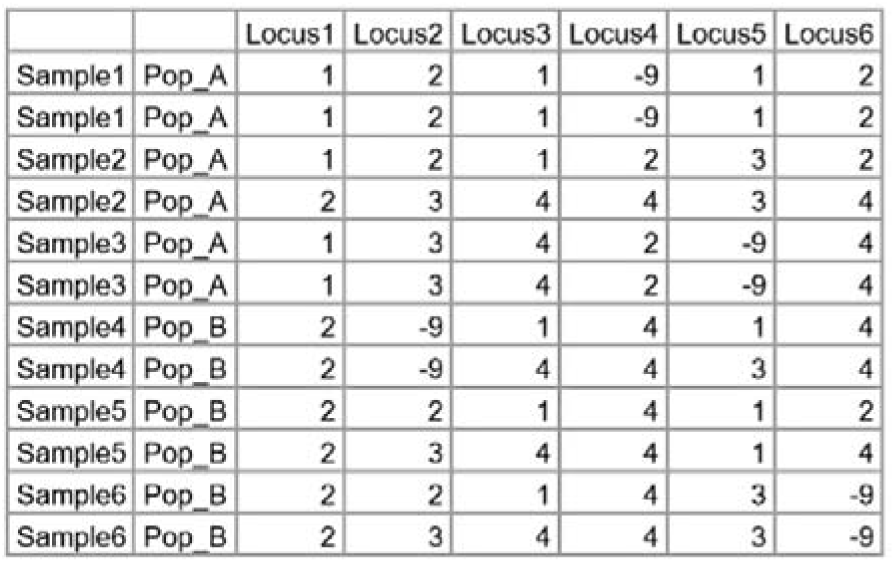
Data example for datasets with two alleles of the same locus are in two consecutive rows.

**S3 Fig.**
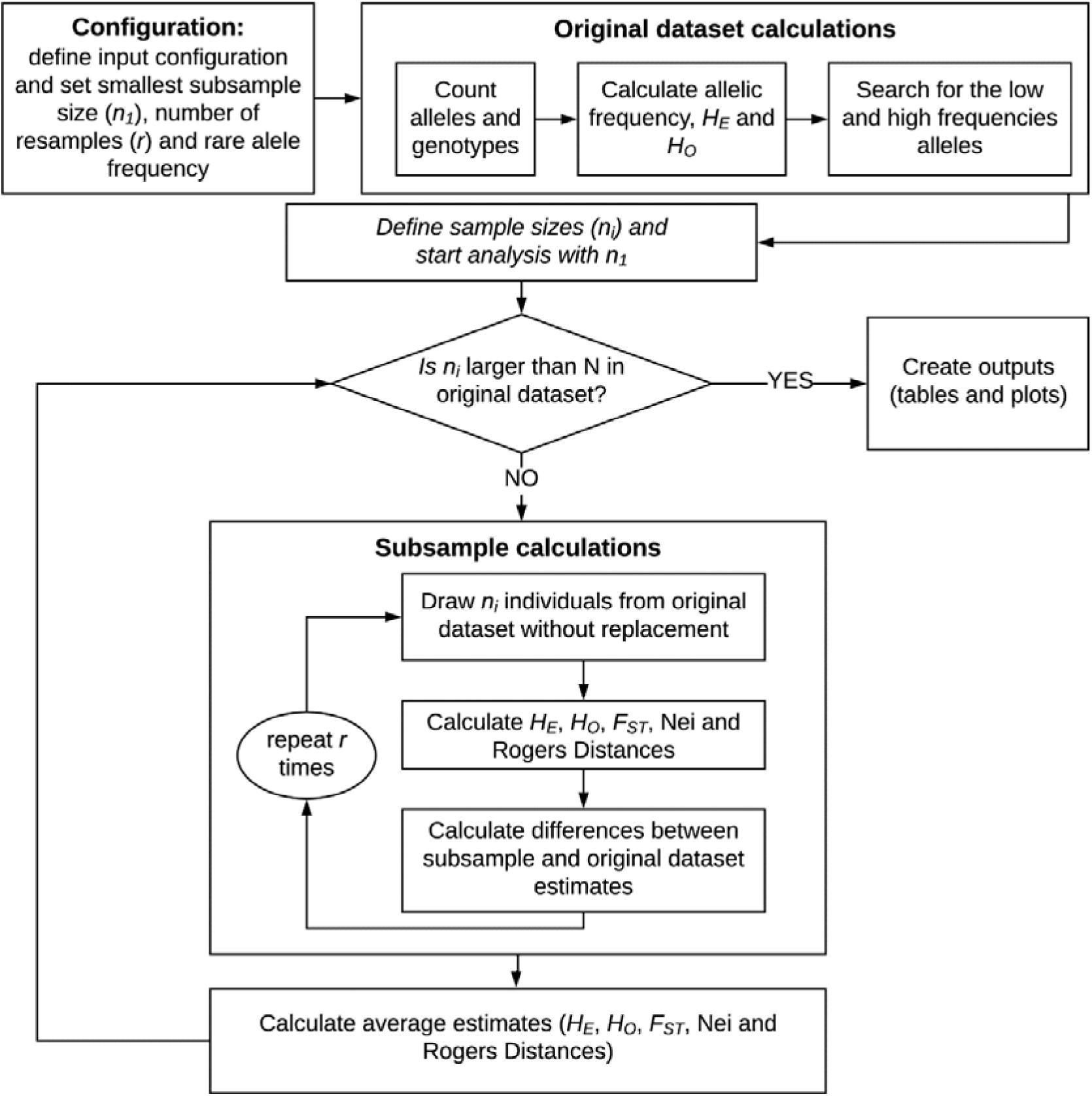
Flowchart of the script.

## Data Accessibility

The script, tutorials, other documentation, and simulated data are available at https://sasii.readthedocs.io/en/latest/index.html. Each empirical dataset availably is described in its corresponding article.

