## Supplementary figures and images for "Sample Size Impact (*SaSii*): an R script for estimating optimal sample sizes in population genetics and population genomics studies"

### S1 Fig

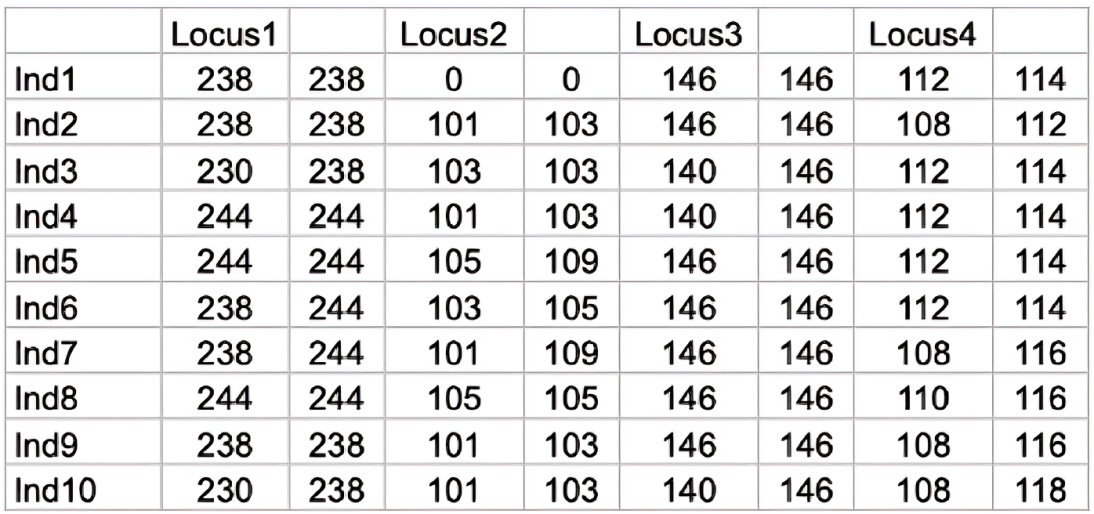

### S2 Fig

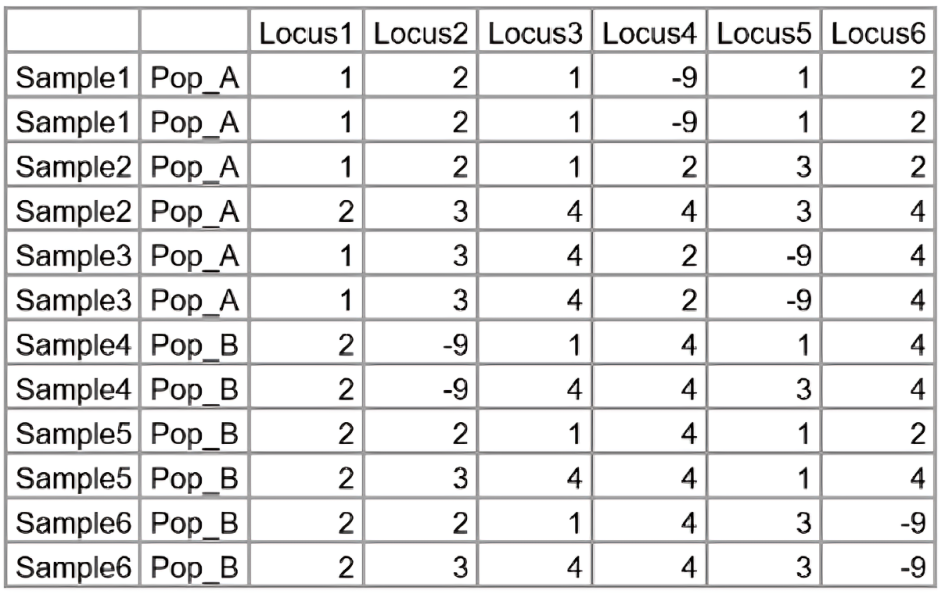

### S3 Fig

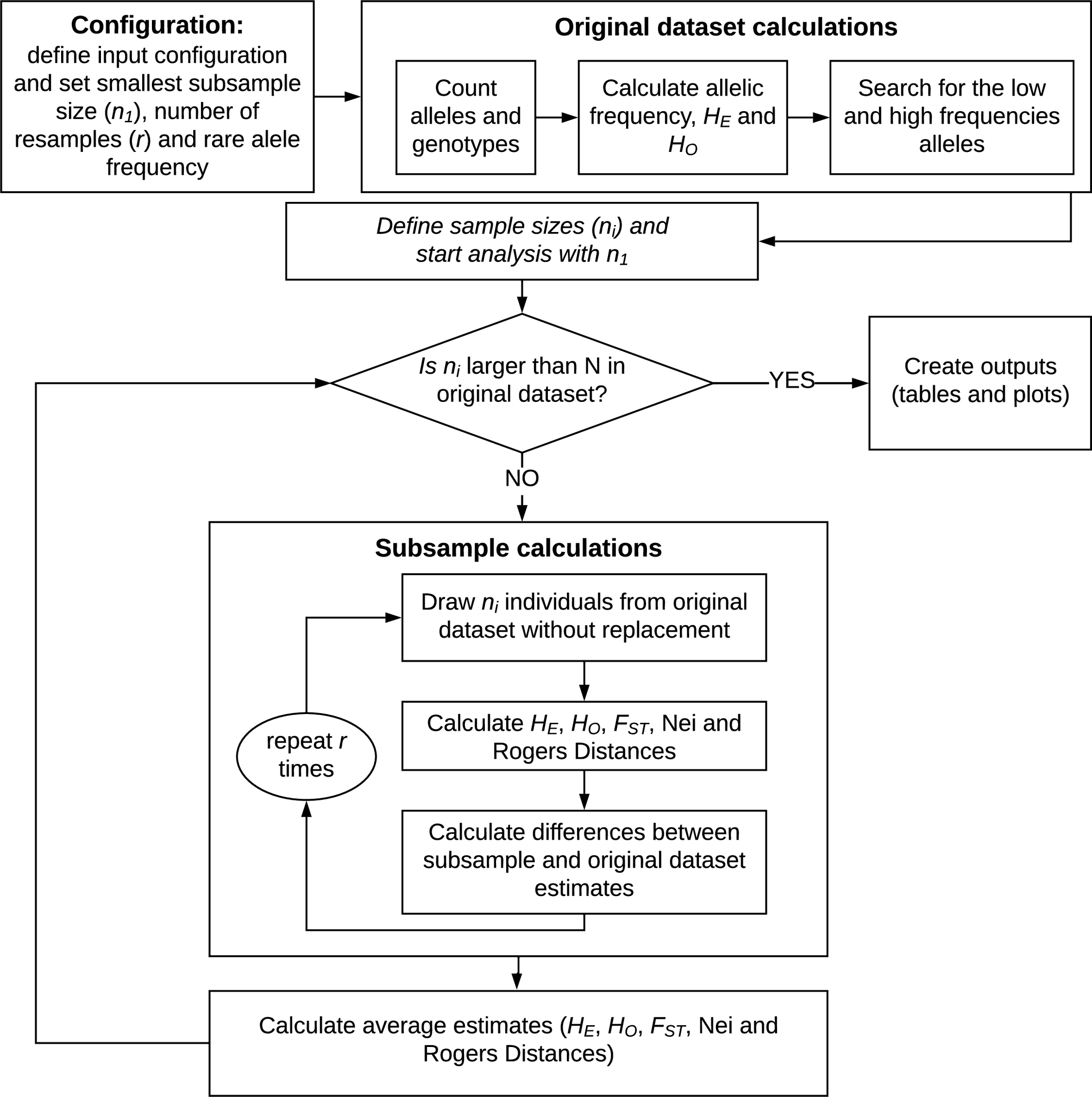
